## Supplementary Figures (1-4) and Information for "Prosit-XL: enhanced cross-linked peptide identification by accurate fragment intensity prediction to study protein-protein interactions and protein structures"

### Supplementary Information 1

**Description of the process for running Percolator in Oktoberfest to rescore CSMs:**

We propose a novel approach in which we use Percolator solely to generate an optimized score for each peptide precursor in a XL-peptide separately by running it on PSM level, rather than on CSM level. This is possible because Prosit-XL generates predictions for each peptide separately and cir-cumvents modeling the four CSM score distributions for the possible true-positive (TP) and false-positive (FP) pairs (TP-TP, TP-FP, FP-TP, and FP-FP) that are present in the target-target (TT, covering TP-TP, TP-FP, FP-TP, and FP-FP), target-decoy (TD, covering TP-FP, FP-TP, and FP-FP) and decoy-decoy (DD, covering FP-FP) matches. When splitting up CSMs into two separate PSMs (one for peptide A and B, each), the clear notion of a target and decoy match remains. Fur-ther, the overall PSM-level score distribution of matches follows the expected behavior as known for linear peptides (Supplementary Fig. S3a). The result of this is a score that is optimized to sepa-rate correct from incorrect PSMs. Because a CSM is incorrect when at least one of the two PSMs is incorrect, we pick the minimum Percolator-optimized PSM-level scores of the two PSMs associ-ated with a CSM as a proxy for the quality of that CSM. This approach should be particularly effec-tive for discriminating lower-scoring TP-TP pairs from higher-scoring TP-FP pairs, since the aggregate score (e.g. by summing) of TP-FP pairs might be overestimated by a good TP candidate. This is often the case in XL-MS, because one of the two peptides is dominating the fragmentation spec-trum. In order to estimate CSM-, peptide pair-, and PPI-level FDR, the new CSM score is passed to xiFDR3 that can model the score distributions of TT, TD, and DD matches correctly.

### Supplementary Information 2

**The list of UL55 cross-link distances for post-fusion (PDBID: 7KDD) and pre-fusion (PDBID: 7KDP):**

Detected cross-links with distance >40 Å for Post-fusion, and <40 Å for Pre-fusion or with distance >40 Å for Pre-fusion, and <40 Å for Post-fusion. The data presented below outlines the distances between specific residues in both the pre-fusion and post-fusion configurations. For each residue, distances are given in the following format:
eg. "Structure, Chain1; Residue1, Chain2; Residue2, Distance"

**Pre-fusion (>40 Å) - Post-fusion (<40 Å)**

1. Post-fusion: 7kdd, Chain A; Residue 1: 568, Chain A; Residue2: 670, Distance Å: 107.4
1. Pre-fusion: 7kdp, Chain A; Residue 1: 568, Chain A; Residue2: 670, Distance Å: 28.7
2. Post-fusion: 7kdd, Chain A; Residue 1: 535, Chain B; Residue2: 670, Distance Å: 77.6
2. Pre-fusion: 7kdp, Chain A; Residue 1: 535, Chain B; Residue2: 670, Distance Å: 32.4
3. Post-fusion: 7kdd, Chain A; Residue 1: 378, Chain B; Residue2: 88, Distance Å: 69.1
3. Pre-fusion: 7kdp, Chain A; Residue 1: 378, Chain B; Residue2: 88, Distance Å: 27.7
4. Post-fusion: 7kdd, Chain A; Residue 1: 670, Chain C; Residue2: 88, Distance Å: 85.4
4. Pre-fusion: 7kdp, Chain A; Residue 1: 670, Chain C; Residue2: 88, Distance Å: 32.1
5. Post-fusion: 7kdd, Chain A; Residue 1: 568, Chain A; Residue2: 691, Distance Å: 135.3
5. Pre-fusion: 7kdp, Chain A; Residue 1: 568, Chain A; Residue2: 691, Distance Å: 32.2
6. Post-fusion: 7kdd, Chain A; Residue 1: 535, Chain C; Residue2: 691, Distance Å: 100.4
6. Pre-fusion: 7kdp, Chain A; Residue 1: 535, Chain C; Residue2: 691, Distance Å: 30.6
7. Post-fusion: 7kdd, Chain A; Residue 1: 568, Chain A; Residue2: 695, Distance Å: 142.6
7. Pre-fusion: 7kdp, Chain A; Residue 1: 568, Chain A; Residue2: 695, Distance Å: 31.4
8. Post-fusion: 7kdd, Chain A; Residue 1: 535, Chain C; Residue2: 695, Distance Å: 108.3
8. Pre-fusion: 7kdp, Chain A; Residue 1: 535, Chain C; Residue2: 695, Distance Å: 36.1
9. Post-fusion: 7kdd, Chain A; Residue 1: 535, Chain A; Residue2: 695, Distance Å: 148.2
9. Pre-fusion: 7kdp, Chain A; Residue 1: 535, Chain A; Residue2: 695, Distance Å: 41.6

**Pre-fusion (<40 Å) - Post-fusion (>40 Å)**

1. Post-fusion: 7kdd, Chain A; Residue 1: 535, Chain C; Residue2: 695, Distance Å: 21.8
1. Pre-fusion: 7kdp, Chain A; Residue 1: 535, Chain C; Residue2: 695, Distance Å: 52.8
2. Post-fusion: 7kdd, Chain A; Residue 1: 209, Chain C; Residue2: 700, Distance Å: No info
2. Pre-fusion: 7kdp, Chain A; Residue 1: 209, Chain C; Residue2: 700, Distance Å: 44.6
3. Post-fusion: 7kdd, Chain A; Residue 1: 378, Chain C; Residue2: 670, Distance Å: 26.0
3. Pre-fusion: 7kdp, Chain A; Residue 1: 378, Chain C; Residue2: 670, Distance Å: 58.1

### Supplementary Figure 1

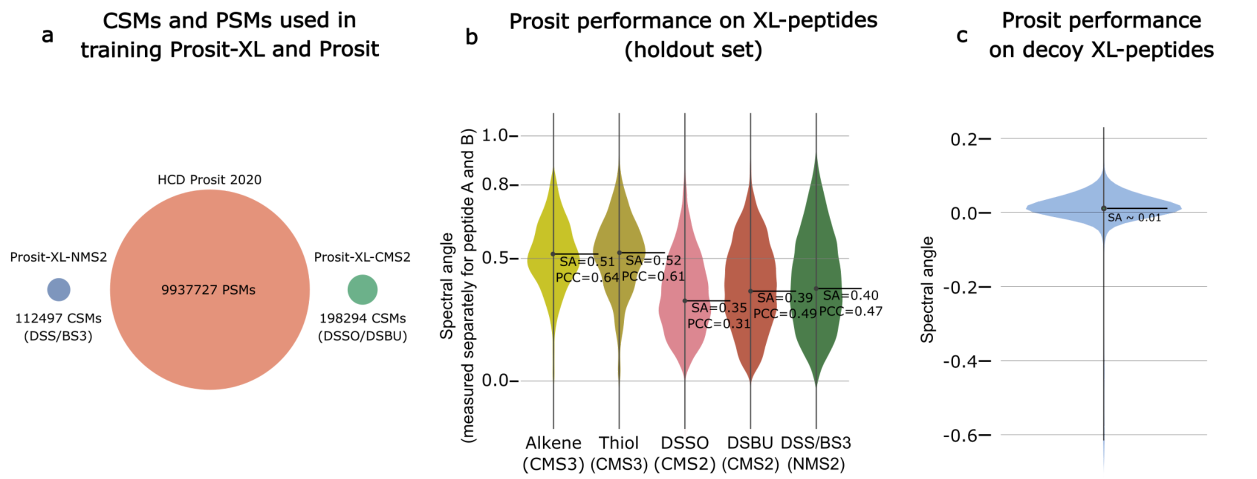


**Supplementary Fig. S1 | Comparison of available training data of Prosit and Prosit-XL and performance evaluation of Prosit on XL-peptides. a)** Comparison of the number of CSMs and PSMs used in training Prosit-XL and Prosit. **b)** Violin plot showing the prediction accuracy of Prosit model for CMS3, CMS2, and NMS2 on the holdout set across 5 different cross-linker types: CMS3-Alkene, CMS3-Thiol, CMS2-DSSO, CMS2-DSBU, and NMS2-DSS/BS3. The black solid line and corresponding numbers indicate the median spectral angle (SA) and Pearson correlation coefficient (PCC). The prediction performance was assessed separately for peptides A and B (PSM level). **c)** Violin plot showing the prediction accuracy of the Prosit model on decoy XL-peptides from the synthetic peptide dataset (DSSO) from replicate one, labeled as 20210203_QExHFX3_RSLC10_DSSO_mainlib_rep1.raw. The measurement was performed exclusively on peptide A.

### Supplementary Figure 2

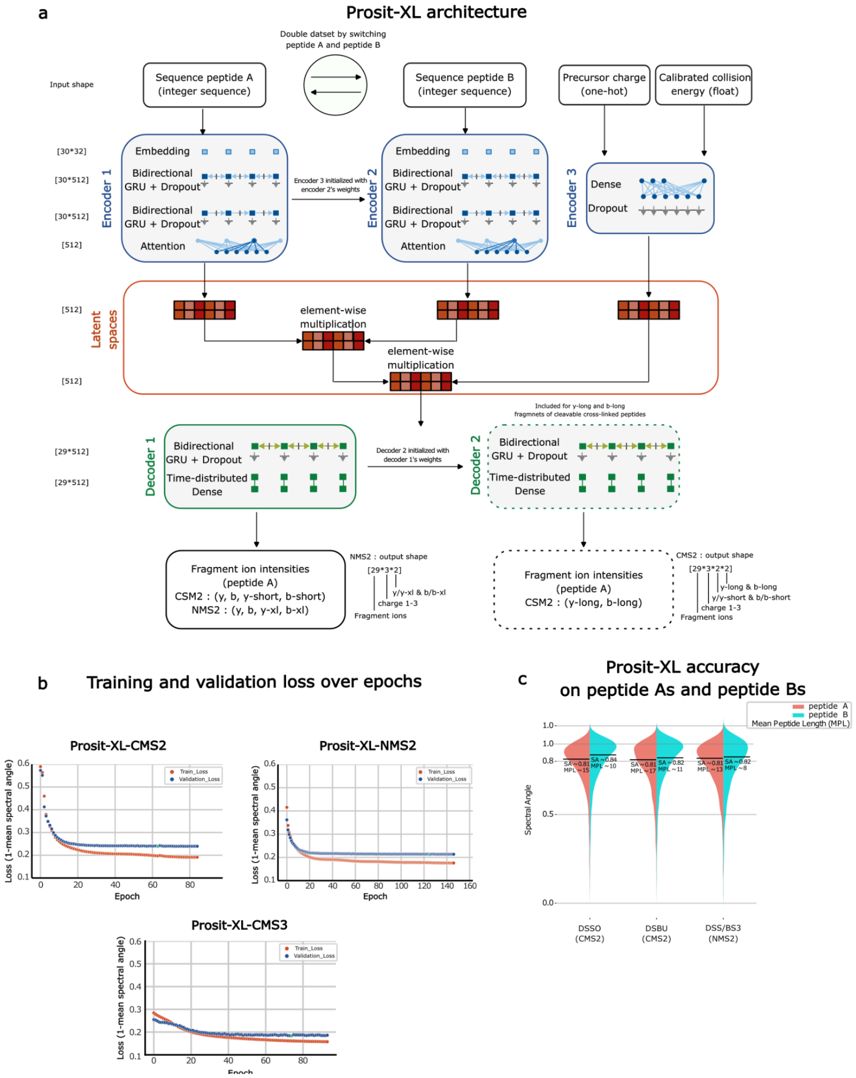


**Supplementary Fig. S2 | Prosit-XL model architecture and training characteristics. a)**. The model takes precursor charge, normalized collision energy and the peptide sequence A and B as input. The encoders for the peptide sequence A and B (encoder 1 and 2) are split in an embedding layer connected to 2 bi-directional recurrent neural networks (BDN) with gated recurrent memory (GRU) units and an attention layer. The encoder 3 consists of one dense layer for precursor charge and normalized collision energy. The encoder 1 and 2 representations are element-wise multiplied and then the resulting output and encoder 3 representations are element-wise multiplied for a fixed size latent space representation. The decoder 1 and 2 for fragment ion intensity prediction consists of one bidirectional GRU resulting in 6 predictions for up to 29 fragmentation positions. The decoder 2 is specifically designed to cover fragments containing the long part of the cleavable crosslinker as modification. **b)** Training and validation loss over epochs. The loss function is calculated by the mean of 1 minus the spectral angle between predicted and experimental fragment ion intensities The orange and blue points show training and validation loss, respectively. **c)** Violin plot comparing the prediction accuracy of Prosit-XL models on peptide As (red) and peptide Bs (light blue) for CMS2 and NMS2 on the holdout set across 3 different cross-linker types: CMS2-DSSO, CMS2-DSBU, and NMS2-DSS/BS3. The black solid line and corresponding numbers indicate the median spectral angle (SA) and the mean of peptide length (MPL).

### Supplementary Figure 3

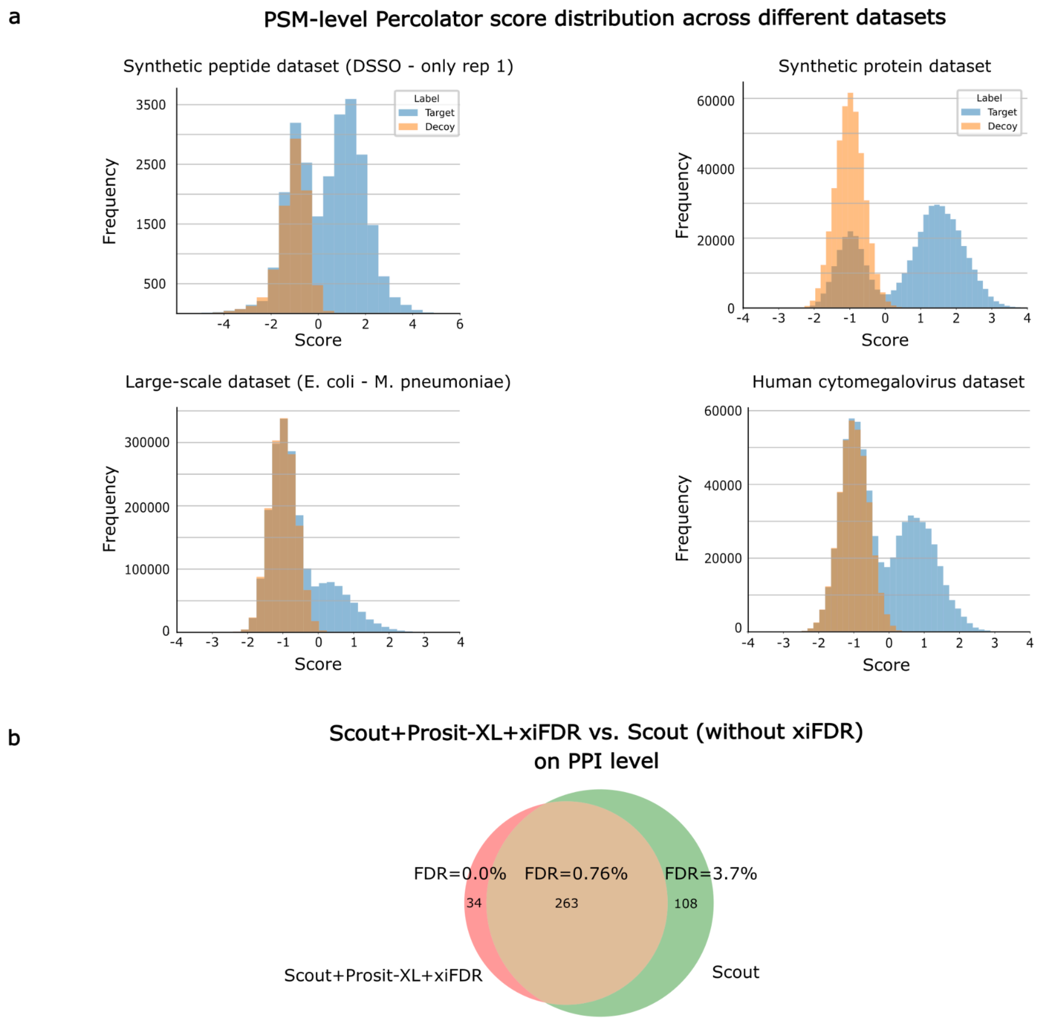


**Supplementary Fig. S3 | Validation of data-driven rescoring approach. a)** PSM-level Percolator score distribution across different datasets including synthetic peptide dataset (upper-left, only rep1), synthetic protein dataset (upper-right), large-scale dataset (*E. coli - M. pneumoniae*, bottom-left) and human cytomegalovirus dataset (bottom-right). Targets and decoys are represented by blue and orange, respectively. **b)** Comparison of Scout+Prosit+XL+xiFDR and Scout (without xiFDR) on PPI level for synthetic protein dataset.

### Supplementary Figure 4

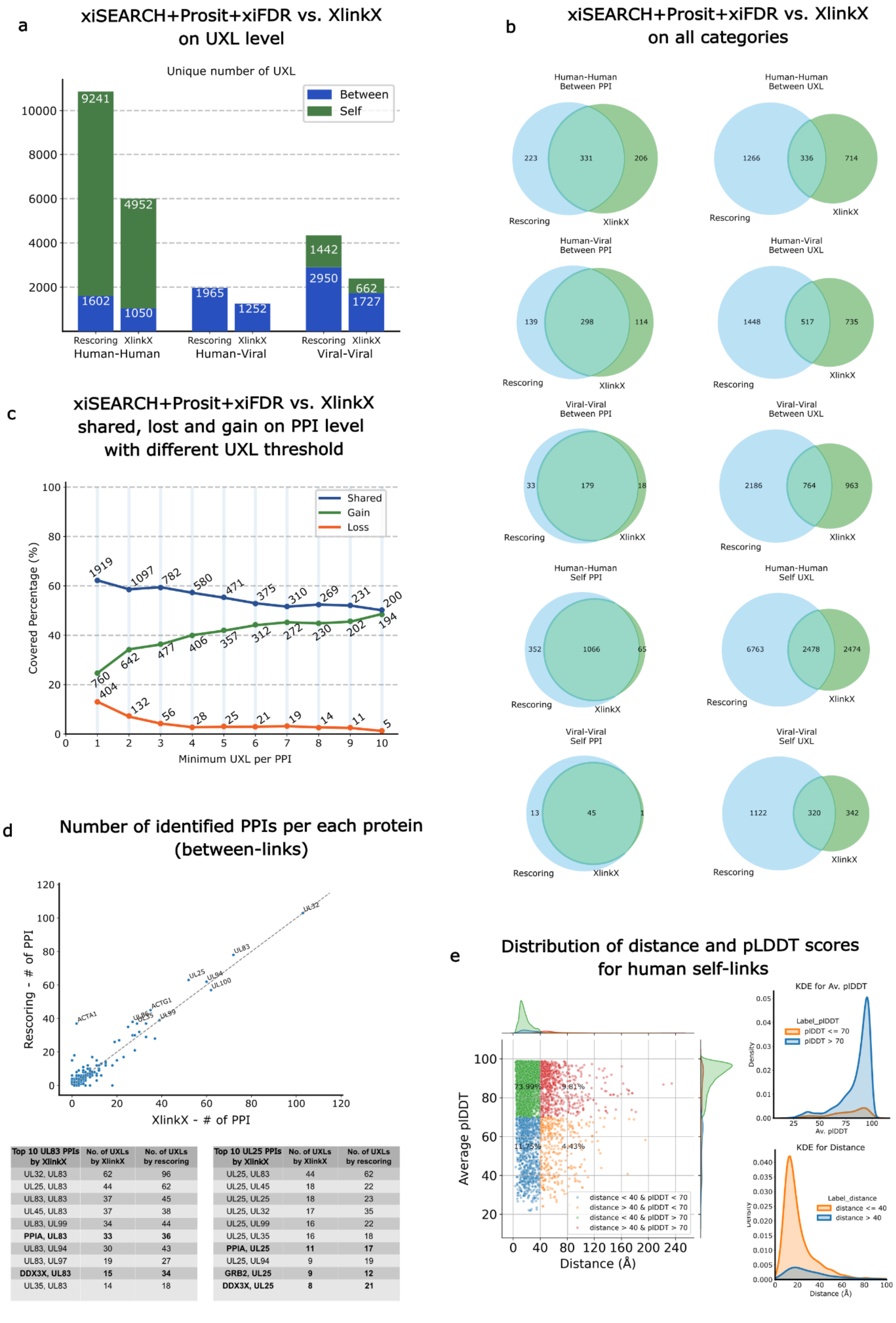


| Top 10 UL83 PPIs by XlinkX | No. of UXLs by XlinkX | No. of UXLs by rescoring |
|---|---|---|
| UL32, UL83 | 62 | 96 |
| UL25, UL83 | 44 | 62 |
| UL83, UL83 | 37 | 45 |
| UL45, UL83 | 37 | 38 |
| UL83, UL99 | 34 | 44 |
| **PPIA, UL83** | **33** | **36** |
| UL83, UL94 | 30 | 43 |
| UL83, UL97 | 19 | 27 |
| **DDX3X, UL83** | **15** | **34** |
| UL35, UL83 | 14 | 18 |

| Top 10 UL25 PPIs by XlinkX | No. of UXLs by XlinkX | No. of UXLs by rescoring |
|---|---|---|
| UL25, UL83 | 44 | 62 |
| UL25, UL45 | 18 | 22 |
| UL25, UL25 | 18 | 23 |
| UL25, UL32 | 17 | 35 |
| UL25, UL99 | 16 | 22 |
| UL25, UL35 | 16 | 18 |
| **PPIA, UL25** | **11** | **17** |
| UL25, UL94 | 9 | 19 |
| **GRB2, UL25** | **9** | **12** |
| **DDX3X, UL25** | **8** | **21** |

**Supplementary Fig. S4 | Evaluation of data-driven rescoring on human cytomegalovirus virion dataset. a)** The bars represent the number of UXLs for human-human, human-viral, and viral-viral interactions identified by xiSEARCH+Prosit-XL+xiFDR (left) and as reported in the original study by XlinkX (right). **b)** Comparison of xiSEARCH+Prosit-XL+xiFDR and XlinkX on both PPI and UXL level for all categories. **c)** Comparison of rescoring and XlinkX at the PPI level, showing shared, gained, and lost interactions when different thresholds of identified UXLs per PPI are applied **d)** The plot represents the number of PPIs per protein (between-links) identified by XlinkX (x-axis) compared to those identified by rescoring (y-axis). The table below lists the top 10 PPIs identified by XlinkX for proteins UL83 and UL25, ranked by the number of UXLs, along with the corresponding UXL counts identified by rescoring **e)** The left plot shows the UXL distances and pIDDT scores of human self-links in four different groups including distance ≤ 40 Å and plDDT ≥ 70 (green points), distance ≤ 40 Å and Av. plDDT < 70 (blue points), distance > 40 Å and Av. plDDT ≥ 70 (red points), and distance > 40 Å and Av. plDDT < 70 (orange points). The top right plot represents the kernel density estimate (KDE) distribution of pIDDT scores for two groups: Av. pIDDT < 70 (orange) and Av. pIDDT > 70 (blue). The bottom right plot represents the KDE distribution of UXL distances for two groups: distance < 40 (orange) and distance > 40 (blue).
